# Dynamics of microbial contaminants is driven by selection during ethanol production

**DOI:** 10.1101/489500

**Authors:** Luciano Lopes Queiroz, Maria Silveira Costa, Alcilene de Abreu Pereira, Marcelo de Paula Avila, Patrícia Silva Costa, Andréa Maria Amaral Nascimento, Gustavo Augusto Lacorte

**Author notes:** Corresponding author: Gustavo Augusto Lacorte.

## Abstract

Brazil is the second largest ethanol producer in the World and largest using sugarcane feedstock. Bacteria contamination is one the most important issues faced by ethanol producers that seek to increase production efficiency. Each step of production is a selection event due to the environmental and biological changes that occur. Therefore, we evaluated the influence of the selection arising from the ethanol production process on diversity and composition of Bacteria. Our objectives were to test two hypothesis, (1) that species richness will decrease during the production process and (2) that Lactic Acid Bacteria will become dominant with the advance of ethanol production due to selection. Bacterial community assemblage was accessed using 16S rRNA gene sequencing from 19 sequential samples. Temperature is of great importance in shaping microbial communities. Species richness increased between the Decanter and Must steps of the process. Low Simpson index values were recorded at the fermentation step, indicating a high dominance of *Lactobacillus*. Interactions between *Lactobacillus* and yeast may be impairing the efficiency of industrial ethanol production.

## 1. Introduction

Ethanol is produced in industries worldwide by the microbial fermentation of the sugars. The yeast *Saccharomyces cerevisiae* is the main species used in ethanol industries, due to their ability of converting carbohydrates into ethanol, energy, cellular biomass, CO_2_, and other products [1]. Brazil is the second largest ethanol producer in the world, and has sugarcanes as the main feedstock [2]. Since 1970, the use of ethanol as fuel for automotive vehicles has increased in Brazil, and decreased in 1990’s due the reduction of subsidies aimed at ethanol industries. A new increase was recorded in the years 2000’s with the advance of flex-fuel cars. Today most cars circulating in Brazil are flex-fuel [3].

There are two types of distilleries used in ethanol production plants in Brazil, fed-batch and continuous fermentation processes. At fed-batch fermentations, tanks are filled one at a time, and tank management and cleaning occur individually, as discrete events. However, in continuous fermentations all tanks are connected in a row and the process occurs simultaneously. The sugarcane juice, after undergoing consecutive treatments steps, is used as substrate for yeast growth and ethanol production [4]. In general, ethanol production consists of cane preparation, juice extraction, juice clarification, juice evaporation, must production, fermentation using yeasts and wine production. The wine follows to other distillation steps and yeast recycling [1].

Maintaining the equipment used in ethanol production within the industry clean is very important to avoid contamination by other microorganisms. Despite the caution in avoiding contamination, it happens and is tolerated [4]. Sugarcane juice has its own associated microbiome and, these microorganisms will be present in some steps of ethanol production. In addition, the soil from sugarcane roots also an important contamination source [1]. Microbial contaminant species may interfere directly in ethanol production competing with the yeast for nutrients during the fermentation process, producing components that may be toxic to the yeast (e.g., acetic and lactic acid)[5], and reducing the carbohydrate consumption and its conversion to ethanol by the yeast. Consequently, high costs in clean up the equipment arise, reducing the productivity in ethanol production [4–6]. The main contaminant taxon is a *Lactobacillus*, specifically a Lactic Acid Bacteria (LAB) found in high proportion during the fermentation procedure [7, 8].

Microbial communities are shaped by four main process: selection, ecological drift, dispersal and speciation [9] (or mutation [10]). Selection is the ability of species to survive and reproduce in different habitats influenced by physical, chemical and biotic factors. Ecological drift depicts the frequencies of species at a given location as a consequence of demographic fluctuations changes. Dispersal is the ability of species to reach and colonize a new location, and, speciation is the origin of new species by speciation events [9, 10]. We believe that the microorganism community structure in the ethanol production process will be determined by selection and ecological drift due the spatial isolation among production steps, which hinder dispersal. In addition, the time from one ethanol production stage to the next is too short for speciation to occur, even in organisms of high reproduction rates [11]. Therefore, each step of production may be considered a selection event due the environmental and biological changes inherent to each stage. In this study we investigated the influence of selection in diversity and composition of Bacteria during ethanol production process.

Our objectives in this study were test two hypothesis (1) species richness will decrease during the process and (2) Lactic Acid Bacteria will become dominant with advance of the ethanol production by selection. Bacterial community assemblage was accessed using 16S rRNA gene sequencing from 19 samples. Microbial contamination is of great importance in the ethanol plant factory. This study can shed light as to how these microbial communities interact and how species succession occurs with advancing of ethanol production.

## 2. Material and Methods

### 2.1. Sampling and ethanol production process

Samples were aseptically collected from each step of the ethanol production process in August 2016 at the Bambuí Bioenergia SA distillery localized in rural area of Bambuí, state of Minas Gerais. We collected one sample from each production step. All samples were stored at − 20 °C until further processing.

The consecutive steps of ethanol production (Figure 1) are sugarcane input, washing, and preparation, followed by sugarcane mill. A same sugarcane sample will pass through six sequential mills. The sugarcane juice is stored in different tanks in each mill, and is latter mixed in one tank. We collected seven samples in this stage: JuiceI, JuiceII, JuiceIII, JuiceIV, JuiceV, JuiceVI, and Mixed Juice. The juice was mixed from sample JuiceVI to Juice II, one by one, resulting JuiceI and a mixture of Juices (II – VI). JuiceI and the previous mixture of Juices (II – VI) were then mixed resulting in the Mixed Juice (Figure 1 – Mixed juice). After the milling step, mixed juice was sent to a heat exchanger (increasing temperature up to ≈50 °C), where liming was conducted. The liming process was carried out to regulate the pH of the solution to 6.2-6.6. The limed juice passed through two other heater exchange processes simultaneously, reaching the temperature of ≈ 105 °C. In this stage we collected the samples HeaterI and HeaterII.

**Figure 1.**
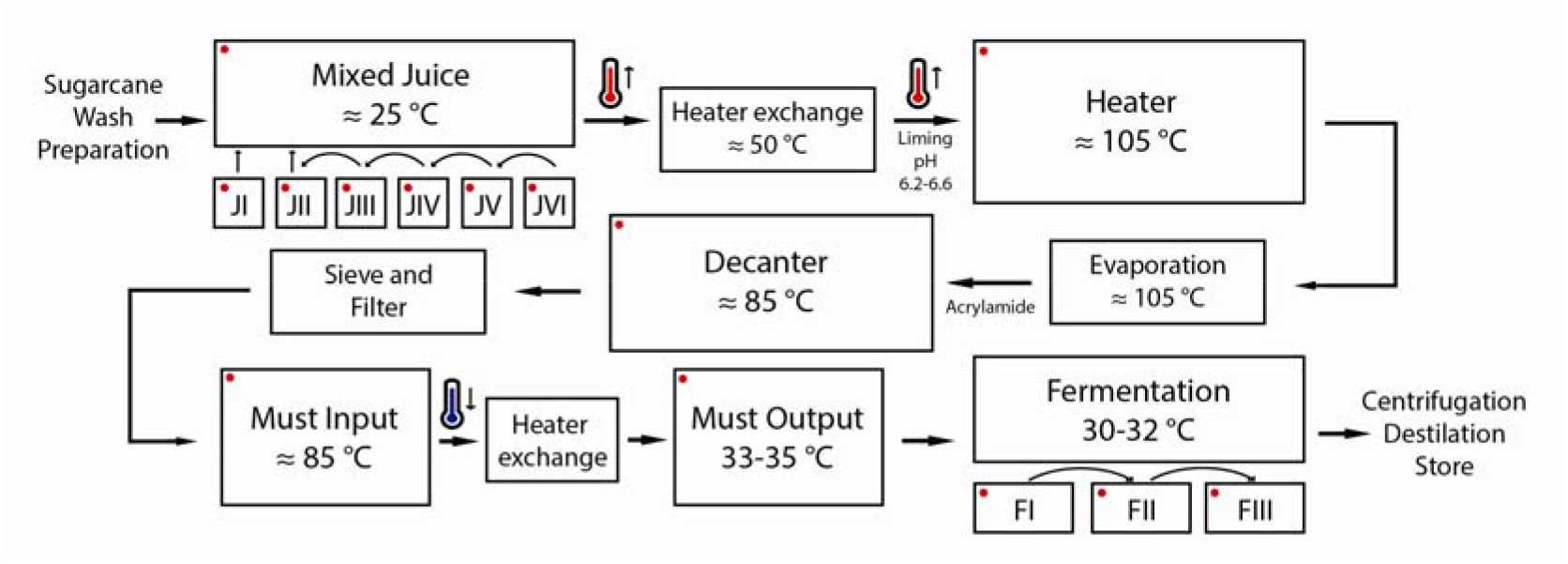
Arrangement of the main ethanol production steps. Red dots show the steps sampled.

The high-temperature juice was sent to the evaporation step, where the non-condensable gases were removed from the juice and acrylamide was added as it promotes the agglomeration of flakes, increases sedimentation speed, etc. These treatments improve the efficiency of the next step – decantation. We collected four decantation samples, two before the material entered the decanter, and two after (DecanterI input, DecanterII input, DecanterI output, and Decanter II output). After decantation, the product was sifted and filtered, and the must (juice treated and evaporated) was sent to the heat exchanger to reduce temperature from ≈ 85 °C to 33-35 °C, temperature needed for yeast growth in fermentation tanks. We collected samples before and after the must passed through the heat exchanger (MustI input, MustI output and MustII output). We were unable to collect must samples before it entered the heat exchanger II.

The low-temperature must was transferred to the first fermentation tank which was filled with yeast solution to up to 25% of its volume capacity. The volume of the tanks is then completed with must (3-4 hours), and the fermentation process takes place for the next 4-5 hours. The must runs through three sequential fermentation tanks, each of which were sampled: FermentationI, FermentationII and FermantationIII.

### 2.2. DNA extraction, library construction and sequencing

Total DNA was extracted from 250 μl of homogenized juice using an E.Z.N.A.^®^ Soil DNA Kit (Omega Bio-tek, Inc., Norcross, GA, USA) following the manufacturer’s instructions with adaptations. DNA was quantified by measuring the absorbance of the sample at 260 nm using a NanoDrop ND-1000 spectrophotometer (Thermo Scientific, Waltham, MA, EUA) and Qubit^®^ dsDNA HS (High Sensitivity) Assay (Life Tecnologies).

The primer set 341F (5’-CCTACGGGNGGCWGCAG-3’) and 785R (5’-GACTACHVGG-GTATCTAATCC-3’) [12] was used to amplify the hypervariable regions V3 and V4 of gene 16S. The Illumina adapter was used to build the 16S sequence library following the protocol provided by Illumina (Illumina, 2013). PCR mixtures contained 0.5 μM each primer, 0.7U of Taq DNA Polimerase (Life Tecnologies, Carlbad, CA., EUA), 1X Buffer, 4mM MgCl_2_, 0.2 mM dNTP, 0.3 mg/ml BSA (Bovie Serum Albumin) and 20 ng of the DNA template. 16S rDNA amplification began with a denaturation time of 3 min at 95 °C, followed by 35 cycles of 45 s denaturing at 95 °C, 1 min primer annealing at 57 °C, 45 s extension at 72 °C and final extension of 10 min at 72 °C. Amplifications were conducted in a Thermal cycler GeneAmp PCR System 9700 (Applied Biosystems, Foster City, CA). All 19 samples were sequenced on the MiSeq platform (Illumina, Inc., USA), using 2□×□250 bp MiSeq V2 reagent kit.

### 2.4. Bioinformatics

We performed a bioinformatics analysis using the Brazilian Microbiome Project (BMP) pipeline [13]. BMP pipeline is a combination of the VSEARCH [14] and QIIME [15] software. The software VSEARCH was used to remove barcodes and primer sequences from a *fastq* file, filter sequences by length (fastq_trunclen 400) and quality (fastq_maxee 0.5), sort by abundance and remove singletons. After that, OTU was clustered and chimeras were removed. Two files were generated through a *fastq* file with filtered sequences and an OTU table in.txt. We assigned the taxonomy to OTU using the *uclust* method in QIIME version 1.9.0 and the SILVA 16S Database (version n128) as reference sequences [16]. The OTU table file was converted to BIOM and taxonomy metadata was added. Sequences were aligned, filtered and the phylogenetic tree constructed. Then, alpha and beta diversity were calculated using QIIME commands. The nucleotide sequence data reported are available in the NCBI under BioProject PRJNA508648.

### 2.5. Statistical analyses

The mean Observed richness, Shannon, Simpson and Equitability values were compared among sampling groups using the Wilcoxon-Mann Whitney test [17]. A heatmap was built with the 20 most abundant OTUs using Ward’s hierarchical clustering method (ward.d2) [18]. Moreover, we used a Principal Coordinates Analysis (PCoA) to compare similarities among samples and tested differences using a Permutational analysis of variance (PERMANOVA) [19]. All analyses were carried out using the statistical software R [20] (*qiimer, ggplot2* [21], *phyloseq* [22], and *vegan* [23] packages).

## 3. Results

### 3.1. Bacterial community diversity

A total of 51.047 sequences and 222 OTU_0.03_ were obtained after bioinformatics analysis. The number of sequences in each sample had a high variability, ranging from 951 sequences in SugarV to 11.159 in MustII-output (mean=2.687±2.338). The last step of the fermentation process had the highest richness (100 species), followed by MustI-input (82), and FermentationI (67). SugarIV was the sample with lowest richness (37). Shannon diversity index (H’) showed values between 0.865 and 2.990. Simpson and Equitability values were similar among samples, considering the similar assumptions of the analysis. Simpson values ranged from 0.288 to 0. 905 and Equitability from 0.238 to 0.679 (Table 1).

**Table 1.**
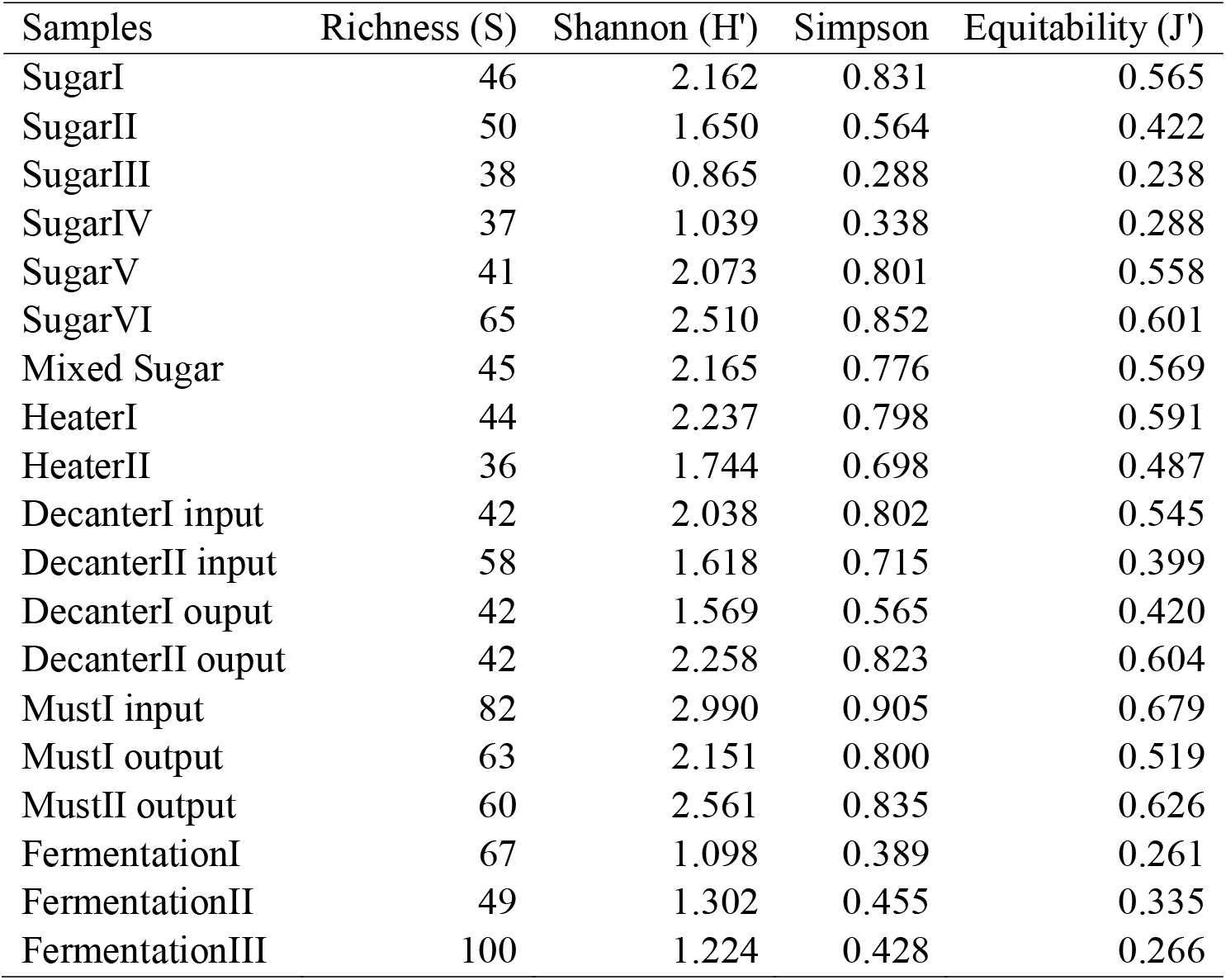
Richness and diversity for in each step of ethanol production.

| Samples | Richness (S) | Shannon (H') | Simpson | Equitability (J') |
| --- | --- | --- | --- | --- |
| SugarI | 46 | 2.162 | 0.831 | 0.565 |
| SugarII | 50 | 1.650 | 0.564 | 0.422 |
| SugarIII | 38 | 0.865 | 0.288 | 0.238 |
| SugarIV | 37 | 1.039 | 0.338 | 0.288 |
| SugarV | 41 | 2.073 | 0.801 | 0.558 |
| SugarVI | 65 | 2.510 | 0.852 | 0.601 |
| Mixed Sugar | 45 | 2.165 | 0.776 | 0.569 |
| HeaterI | 44 | 2.237 | 0.798 | 0.591 |
| HeaterII | 36 | 1.744 | 0.698 | 0.487 |
| DecanterI input | 42 | 2.038 | 0.802 | 0.545 |
| DecanterII input | 58 | 1.618 | 0.715 | 0.399 |
| DecanterI ouput | 42 | 1.569 | 0.565 | 0.420 |
| DecanterII ouput | 42 | 2.258 | 0.823 | 0.604 |
| MustI input | 82 | 2.990 | 0.905 | 0.679 |
| MustI output | 63 | 2.151 | 0.800 | 0.519 |
| MustII output | 60 | 2.561 | 0.835 | 0.626 |
| FermentationI | 67 | 1.098 | 0.389 | 0.261 |
| FermentationII | 49 | 1.302 | 0.455 | 0.335 |
| FermentationIII | 100 | 1.224 | 0.428 | 0.266 |

Species richness was consistent in the first three steps (Sugar, Heater, and Evaporation), but we observed that richness increased in the Must step, followed by a low richness increase during Fermentation. Richness of the Evaporation and Must groups differed (Wilcoxon-Mann Whitney test, *p* = 0.044) (Figure 2). The diversity indices measured (Shannon, Simpson and Equitability) exhibited similar results as regards abundance distribution among samples and groups of ethanol production steps. Diversity index values were high in the Sugar, Heater, Evaporation, and Must steps with mean values of 1.977±0.546 (Shannon), 0.711±0.182 (Simpson), and 0.506±0.123 (Equitability). When must was transferred to the fermentation tanks, these values decreased (mean values 1.208±0.102, 0.423±0.033, and 0.287±0.411, respectively; Suppl. figures 1-3 for diversity indices). These differences were tested by Wilcoxon-Mann Whitney test and no significant difference was found.

**Figure 2.**
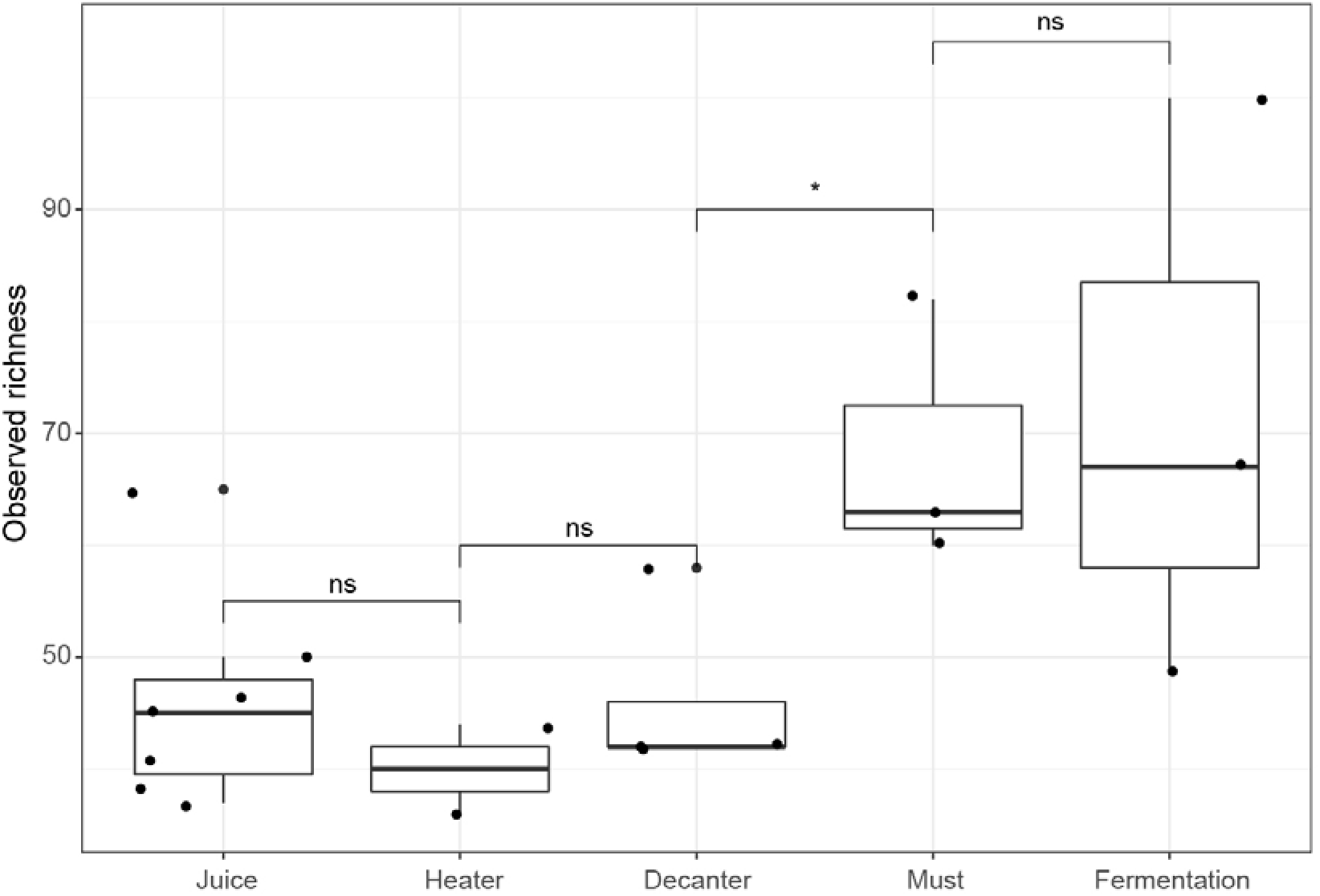
Species observed in each step of ethanol production.

### 3.2. Bacterial community composition

The community composition of all samples was comprised mostly Firmicutes (76.43%), Proteobacteria (19.46%), and Cyanobacteria (3.71%) (Suppl. Figure 4). The Firmicutes phylum was the most abundant in several samples and the sequences from this phylum were in the Bacilli class (75.31%; Suppl. Figure 5). Over 90% of the sequences in all Juice samples were Bacilli, except in JuiceI, where 53.58% of the sequences were Bacilli, 21.83% were Alphaproteobacteria (Proteobacteria) and 13.56% of Chloroplast (Cyanobacteria). Bacilli were highly dominant in Heater samples (on average 94.62% of the sequences), and the mean was 81.45% in Fermentation samples. The relative abundance for the Evaporation and Must samples was a similar to that recorded for JuiceI.

The relative genera abundance showed a great variability among the ethanol production steps (Figure 3). The production steps are linear starting with sugarcane mill and juice production. The first juice obtained was JuiceI and comprised the genera *Weissella* (27.3%), *Leuconostoc* (22.01%), and *Oryza mayeriana* (rice; 19.67%). The abundance of *Lactobacillus* increased slightly from JuiceII to JuiceIV (11.65%), *Weissella* showed a high increase to 79.27%, and *Leuconostoc* a high decrease to 0.55%. JuiceV had a high abundance of Lactobacillus (55.31%), followed by *Weissella* (28.18%) and *Streptococcus* (9.57%). JuiceVI had different proportions of these genera, specifically a *Lactobacillus* reduction to 29.61%, *Weissella* reduction to 8.67% and *Streptococcus* increase to 32.7%. The last step of juice production is mixing all juices in a same tank. The Mixed Juice sample had a high abundance of *Streptococcus* (43.54%), followed by *Lactobacillus* (37.34%) and a high decrease of *Weissella* abundance (0.37%). These patterns may also be observed in the heatmap figure (Figure 4).

**Figure 3.**
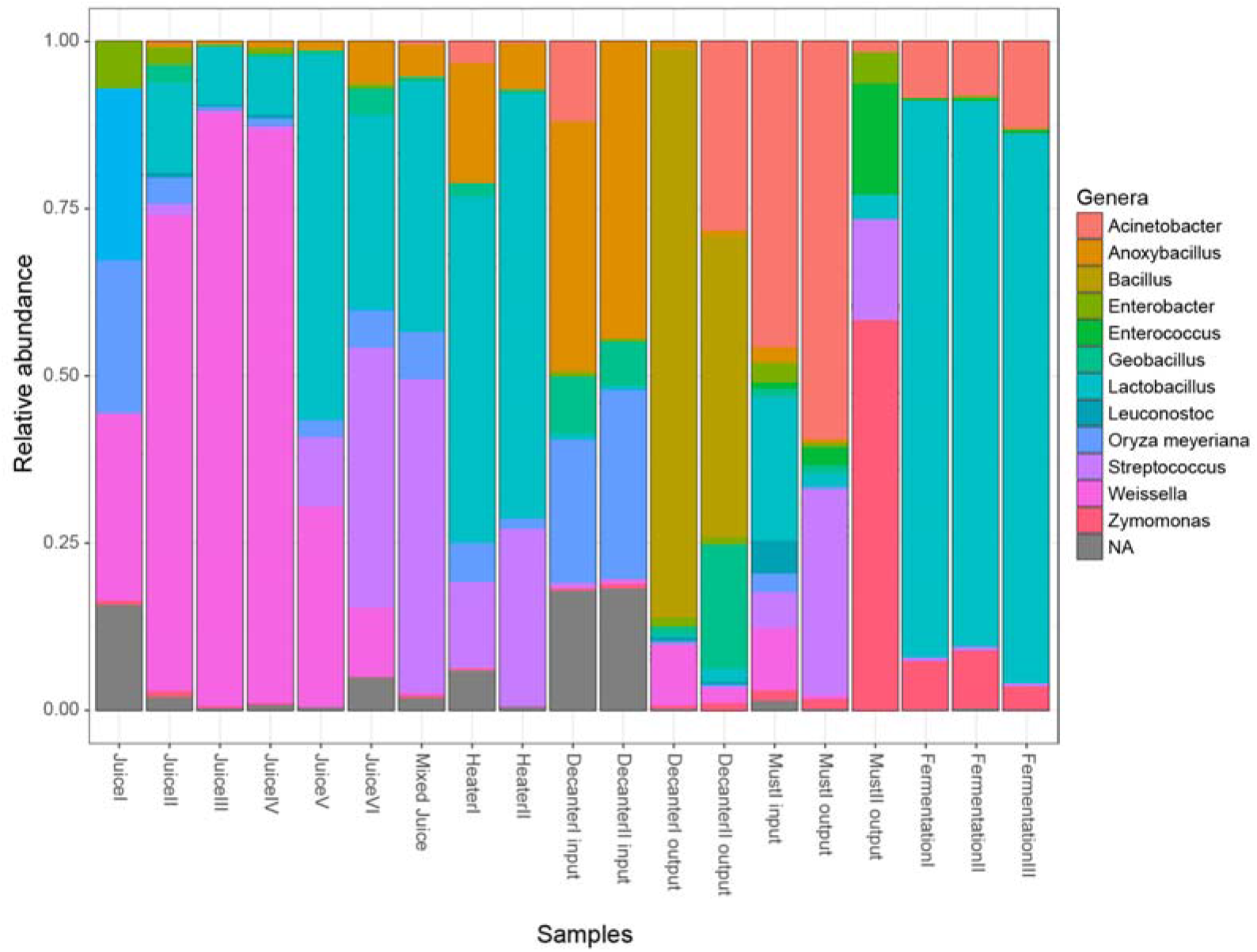
Relative abundance of the main genera found in each step of ethanol production.

**Figure 4.**
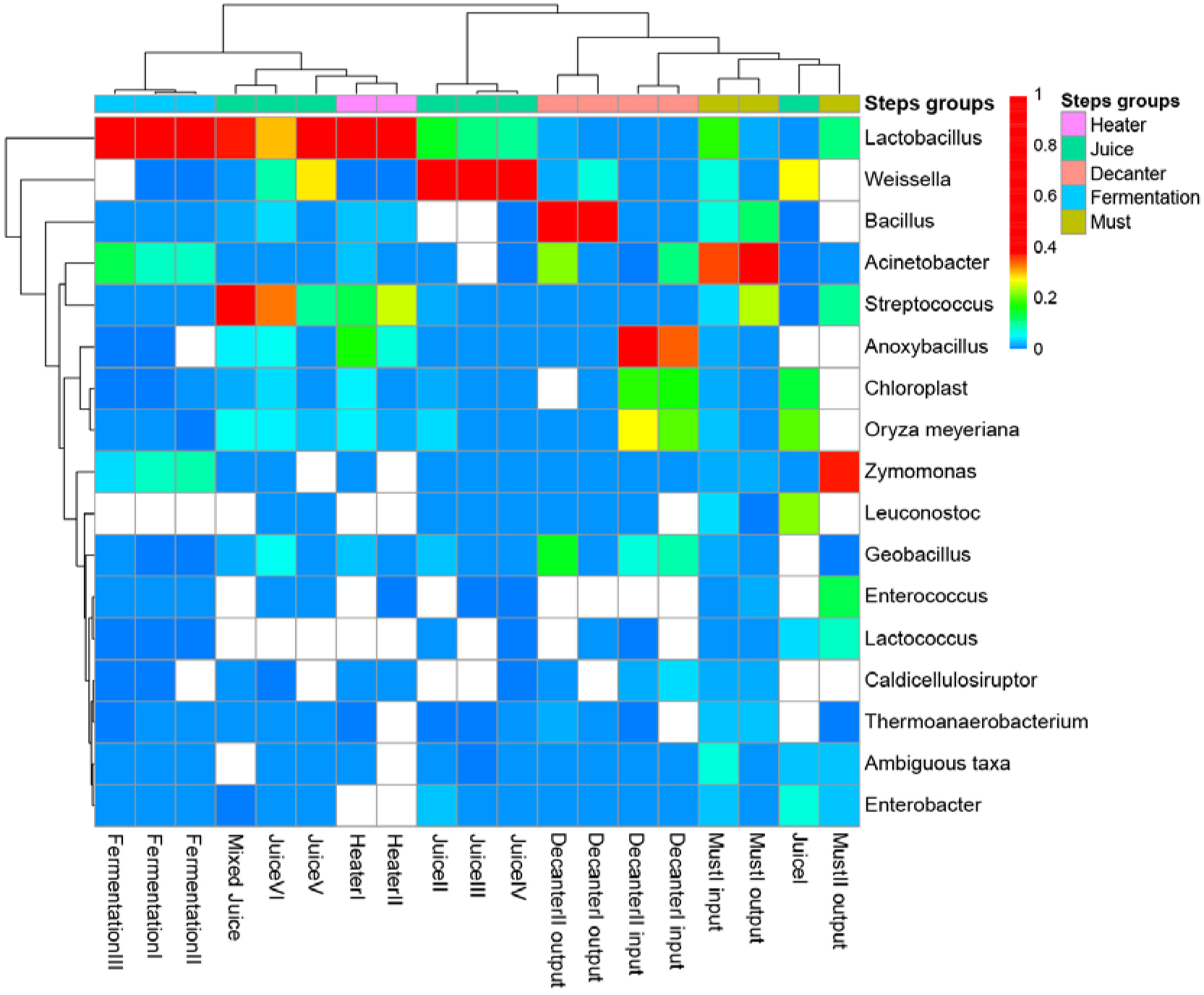
Heatmap of the 20 most abundant OTU0.03 found in each step of ethanol production.

The next step of ethanol production is the Heater step, in which the Mixed Juice passes through two heaters simultaneously (HeaterI and II). The *Lactobacillus* abundance in the juice after it passed through HeaterI was 49.55%, followed by *Anoxybacillus* with 17.03% and Streptococcus with 12.63%. After HeaterII, *Lactobacillus* abundance increased to 61.91%, *Streptococcus* to 24.95% and *Anoxybacillus* decreased to 6.41%. The Juice passed through the heaters was sent to the decantation tank for the evaporation step (two tanks). We collected samples from the evaporation step before juice was directed to the tanks (Evaporation I input and Evaporation II input) and after the end of the evaporation process (Evaporation I output and Evaporation II output). Evaporation I input and Evaporation II input had a similar genera pattern, specifically, *Anoxybacillus* was the most abundant (34.38% and 42.19%, respectively), followed by *Oryza meyeriana* (19.64% and 28.86%, respectively). We observed a dominance of the genus *Bacillus* after evaporation processes, comprising nearly 80% of the sequenced genes Evaporation I output and 50% in Evaporation II output. *Acinetobacter* and *Geobacillus* were also abundant in Evaporation II output sample (21.48 and 14.32%, respectively).

The must produced after evaporation was sent to the heat exchanger to reduced temperature to the reach optimum temperature for the yeast. The most abundant genus in MustI input was *Acinetobacter* (31.44%), followed by Lactobacillus (17.43%) and Streptococcus (4.06%). The pattern changed in MustI output, with an increased *Acinetobacter* (46.43%) and Streptococcus abundance (24.09%) and decreased Lactobacillus abundance (2.01%). MustII output is a mixture of MostI and II output, and showed a high abundance of *Zymomonas* (37.34%), followed by *Lactobacillus* (10.45%), *Streptococcus* (9.71%), and prominent reduction of *Acinetobacter* (0.95%).

The fermentation process occurs in three different and sequential tanks, FermentationI, II and III. *Lactobacillus* was the most abundant genus in all fermentation steps, specifically accounting for 80.39% of the sequences in FermentationI, 77.95% in FermentationII, and 79.83% in FermentationIII, followed by *Acinetobacter* and *Zymomonas*.

A Principal Coordinates Analysis (PCoA) was conducted with four distance matrices: Bray-Cutis (Figure 5), Jaccard, and Weighted and Unweighted UniFrac (Suppl. Figure 6). The samples were sorted into five groups according to the step of production. Microbial communities present in samples from the same group, when using Bray-Curtis similarity matrix, were highly similar (PERMANOVA, F = 4.667, R^2^ = 0.55, *p* = 0.001). We observed a high proximity between samples from Fermentation step (blue dots, figure 5). Juice samples were grouped in left side of the ordination plot (near the - 0.2 position of Axis 1), and Must and Decanter samples were grouped in the center.

**Figure 5.**
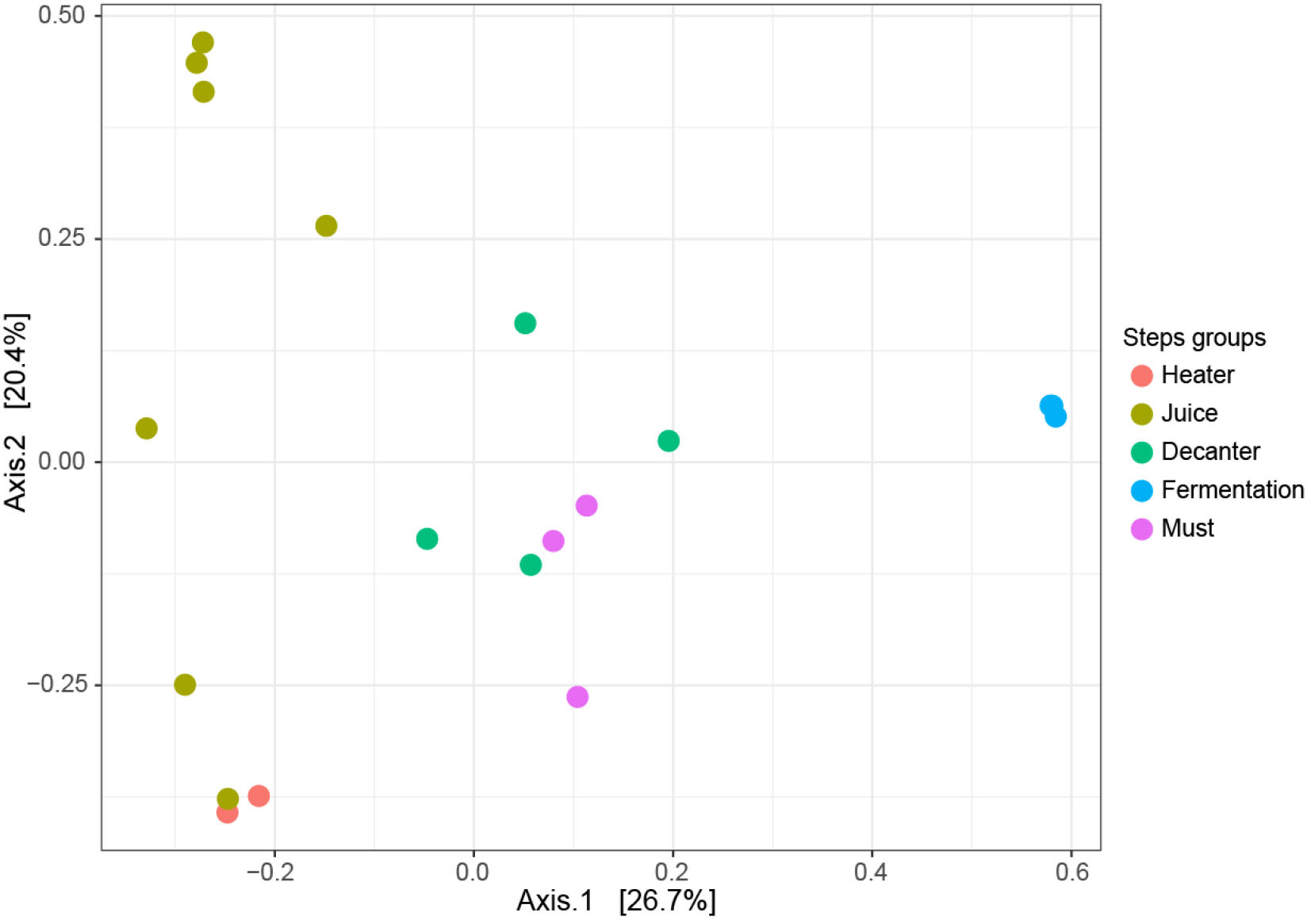
Principal Coordinates Analysis (PCoA) using Bray-Curtis dissimilarity matrix for the samples of ethanol production process.

## 4. Discussion

### 4.1. Bacterial richness increase even after selection events

We observed that bacterial richness increased, instead decreasing as we expected in our first hypothesis. In this study, each step in the ethanol production process was considered a selection event for microbial interactions and/or environmental changes. Bacterial species richness was constant during first three steps (Juice, Heater, and Decanter). However, richness increased from the Decanter to the Must steps (Figure 2). This result may be explained by a contamination with microorganisms after the decantation process.

Industrial ethanol production is not a sterile process, microbial contamination is common and identified as an important problem regarding ethanol production efficiency [4, 24]. The increase of microbial richness after the decantation process, as observed in this study, showed that selection events that occur before decantation may be efficient in controlling microbial communities. The samples may become contaminated during the transfer of decanted juice to must tanks, when the juice passes the rotary sieves, or during the evaporation system.

The species richness in Must samples was constant during the Fermentation step. However, evenness decreased in Fermentation samples, as we observed in the diversity indices values. Relative abundance is altered by selection, such that the environmental and biological condition may be ideal for the survival and reproduction of a species at a given time, and not at another [10]. Diversity indices based on observed richness and relative abundance, such as the Shannon, Simpson, and Equitability indexes, may be used to understand certain species and their contribution in abundance [25, 26].

However, the differences in diversity index values between the Must and Fermentation stages were not significant (Suppl. Figure 1-3).

### 4.2. Selection process shaping microbial community

The high variance of the relative abundance of OTUs among the ethanol production steps may be influenced by temperature [4] and negative interactions with other taxon. The importance of these two variables for selection may be noted when observing the relative abundance of *Weissella. Weissella* exhibited a high abundance in Juice samples. However, its abundance was lower in the juice mixture, possibly due to a negative interaction with *Lactobacillus* and *Streptococcus*. This negative interaction may have reduced *Weissella* while increasing *Lactobacillus* and *Streptococcus* abundance values. The sequences rated as *Weissella* almost disappeared after the Heater steps (Figure 3). A recent study found lower abundances of *Weissella* in fermentation tanks,[8] as recorded in this study.

All of our sequences of *Weissella* were rated at species level as *Weissella confuse. Weissella* species are non-spore and gram-positive coccobacilli [27]. *W. confuse* is found in fermentation foods, but have also been associated to human diseases (sepsis and bacteremia) as an opportunistic pathogen [28]. These bacteria are heterofermentative, belong to the Lactic Acid Bacteria (LAB) group, some strain are able to growth at 45 °C and consume several types of carbohydrates (Fructose, Galactose, Maltose) [29]. *W. confuse* has been found in sugarcane samples [24, 30]. The reduced abundance of *Weissella* may be explained by a low ability to uptake carbohydrates from the environment when compared with other lactic acid bacteria, such as *Lactobacillus* and *Streptococcus*.

On the next steps, temperature increased twice to approximately 50 °C and to 105 °C. We observed that *Lactobacillus* and *Streptococcus* were the most abundant species in Heater samples, even under high temperatures. The relative abundance of *Lactobacillus* and *Streptococcus* decreased abruptly, while the abundance of *Anoxybacillus* and *Geobacillus* increased, occupying the habitat when juice samples are stored in decanter tanks, after undergoing the acrylamide treatment and being kept under constant temperature at 105 °C (Decanter input samples).

*Anoxybacillus* and *Geobacillus* are gram-positive Bacillaceae family, and have the morphology of rod or coccus [31, 32]. Both are moderately thermophilic, with optimal growth temperature of 50-62 °C and 55-65 °C for *Anoxybacillus* and *Geobacillus* respectivelly. These genera are found in several types of habitats, for example geothermal springs, manure, and milk-processing plants. An interesting observation about these two genera is their ability to form endospores. *Geobacillus* have a widespread distribution in world, even in cold habitats [33]. *Anoxybacillus spp*. are alkaliphilic or alkalitolerant, but are able to grow under neutral pH [34]. The high abundance of *Anoxybacillus* and *Geobacillus* in the decantation step may be explained by their capacity of tolerating high temperatures, unlike others species. Thus, they dominated habitat due to the high temperature and absence of competitors.

The juice passed through rotary sieves and another evaporation system during the transportation of the juice from Decanter to Must tanks. The temperature of the decanted juice before entering Must tanks ranged from 85-90 °C, and its microbial communities were dominated by *Bacillus*. This temperature reduction was enough to create conditions for *Acinetobacter* growth, as confirmed on the next step with a high increase of its relative abundance. *Acinetobacter* is gram-negative Protobacteria and has been found in soil environments [35]. These bacteria may have come with sugarcane roots, despite the washing steps.

When the temperature of the MustI output sample was reduced to 33-35 °C, the abundance of *Acinetobacter* remained high, but abundance of *Streptococcus* began to increase. This sample was mixed with MustII output, and presented a high increase of *Zymomonas* abundance and decrease of *Acinetobacter* under mesophilic conditions. The genus *Zymomonas* has only one species, *Zymomonas mobilis*. *Z. mobilis* is a gram-negative Alphaproteobacteria, facultative anaerobic and rod-shaped, it can produce bioethanol by the Entner–Doudoroff pathway [36]. This species has been studied for industrial ethanol production, and it has advantages over *Saccharomyces cerevisiae*, such as the high ethanol tolerance [37]. The high abundance at this moment may be explained by optimal temperature conditions and the absence of other fermentative microorganisms in high abundance. *Lactobacillus* became dominant and *Zymomonas* abundance reduced to a stable proportion when Must was sent to fermentation tanks and yeast was added.

We also observed a high similarity in the composition of the microbial communities within samples from the same group (i.e., fermentation step; Figure 5). Samples from Juice group had low within-group similarity values (yellow-mustard dots). However, when Mixed Juice was submitted to the first step of juice preparation (HeaterI and II), the communities present became highly similar (red dots). Therefore, communities were selected by the increased temperature and pH regulation with liming. These results may be explained by the different environmental variables and inter-specific interactions present at each community, in each production step.

### 4.3. High abundance of Lactobacillus during fermentation

Our second hypothesis was that Lactic Acid Bacteria (LAB) would become with the advance of ethanol production, due to selection. Lactobacillus was the most abundant bacteria in the Fermentation stage. Other LAB, such as *Weissella*, showed a lower abundance and *Leuconostoc* was not present at this ethanol production step. Several studies have recorded *Lactobacillus* as dominant species during the fermentation of sugarcane juice. Costa et al. (2015) reported that almost 100% of the sequences were of *Lactobacillus* species, while Bonatelli et al. (2017) reported a prevalence of 92 to 99%.

The main selection factors present during the fermentation step were temperature (30-32 °C), high abundance of yeast competing for carbohydrates, and ethanol produced during carbohydrate fermentation [4, 7]. Ethanol is a toxic compound for most of the microorganisms present in sugarcane juice during fermentation [24, 38]. *Lactobacillus*, *Zymomonas, Acinetobacter, Enterococcus* and *Weissella* were resistant, but *Lactobacillus* was a better competitor in carbohydrate uptake than the others and became dominant because of its fast growth [39].

### 4.4. Conclusions and remarks

Microbial communities varied greatly among the ethanol production process, with main changes occurring due to the influence of temperature. Thus, temperature was identified as an important selection factor. Our first hypothesis was rejected, in contrast with our expectations, given that species richness increased between Decanter and Must samples. Our second hypothesis was corroborated by the compositional data, which show a high abundance of *Lactobacillus* sequences in the Fermentation step. However, no significant differences were recorded among the diversity indices of the production steps. The effort made to avoid contamination in ethanol industries and the identification of temperature as important selection factor, is not enough to avoid the prevalence of *Lactobacillus*, a major competitor with yeast for resources in this system.

## Supporting information

## 5. Acknowledges

We would like to thank Bruno Spacek for critical reviews of the manuscript.

