## Supplementary material for "Dynamics of microbial contaminants is driven by selection during ethanol production"

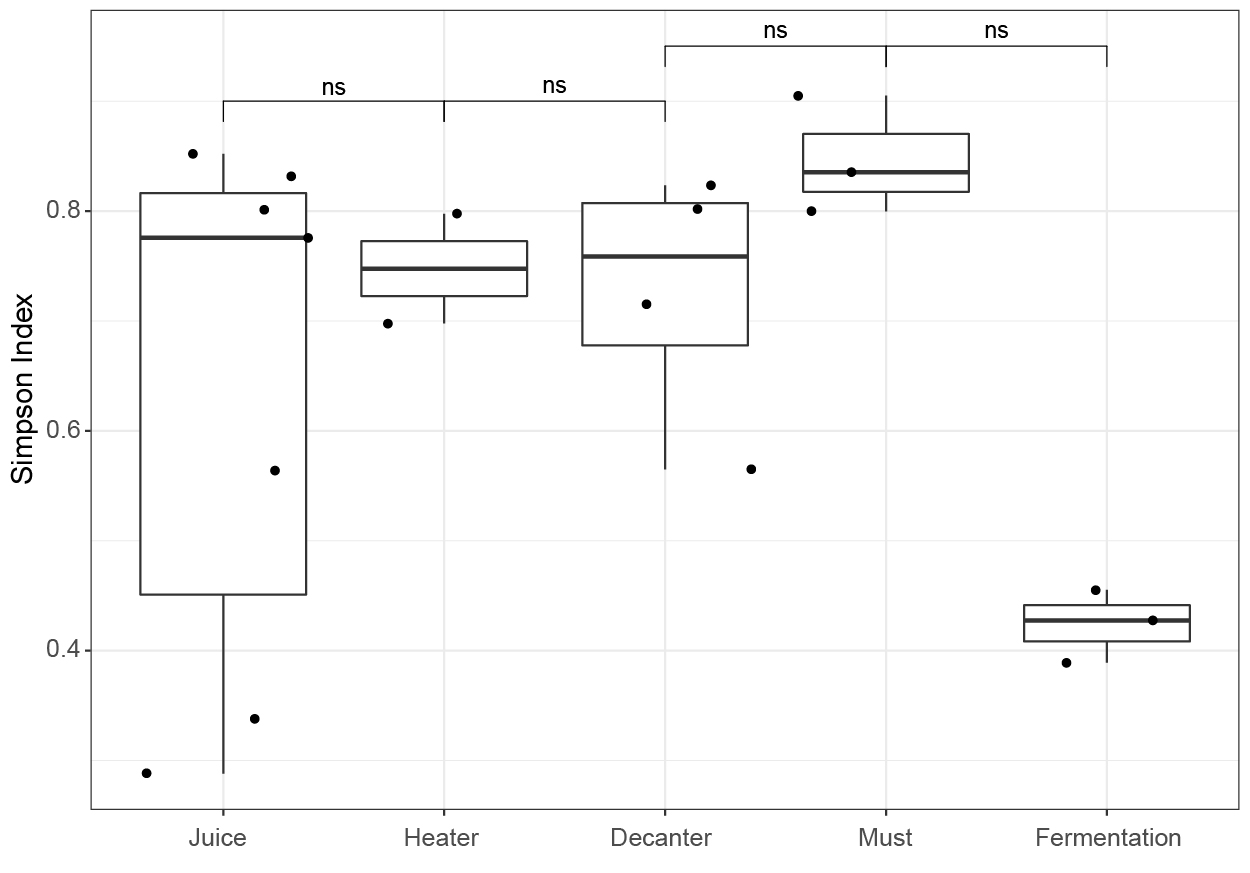

**Supplementary figure 1:** Simpson’s index in each step of ethanol production.

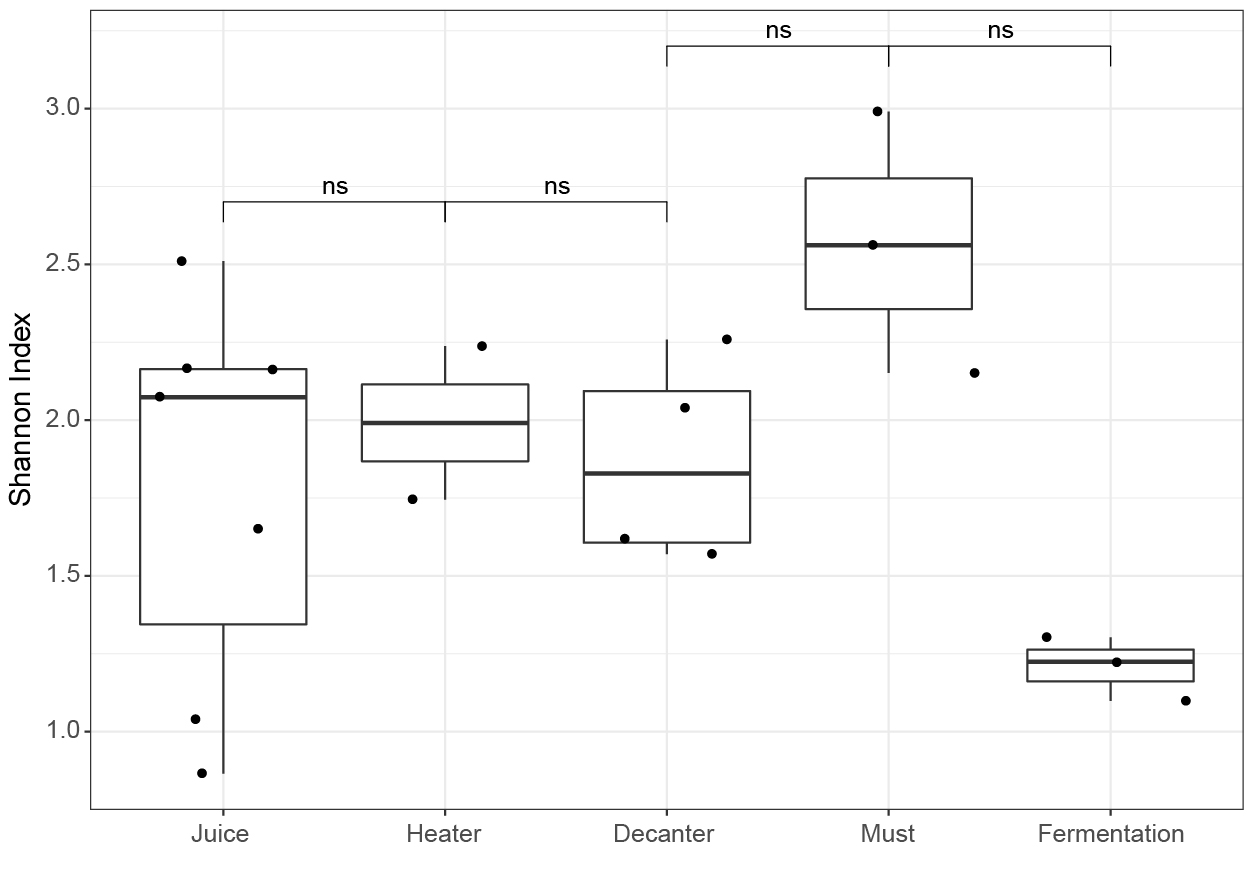

**Supplementary figure 2:** Shannon index in each step of ethanol production.

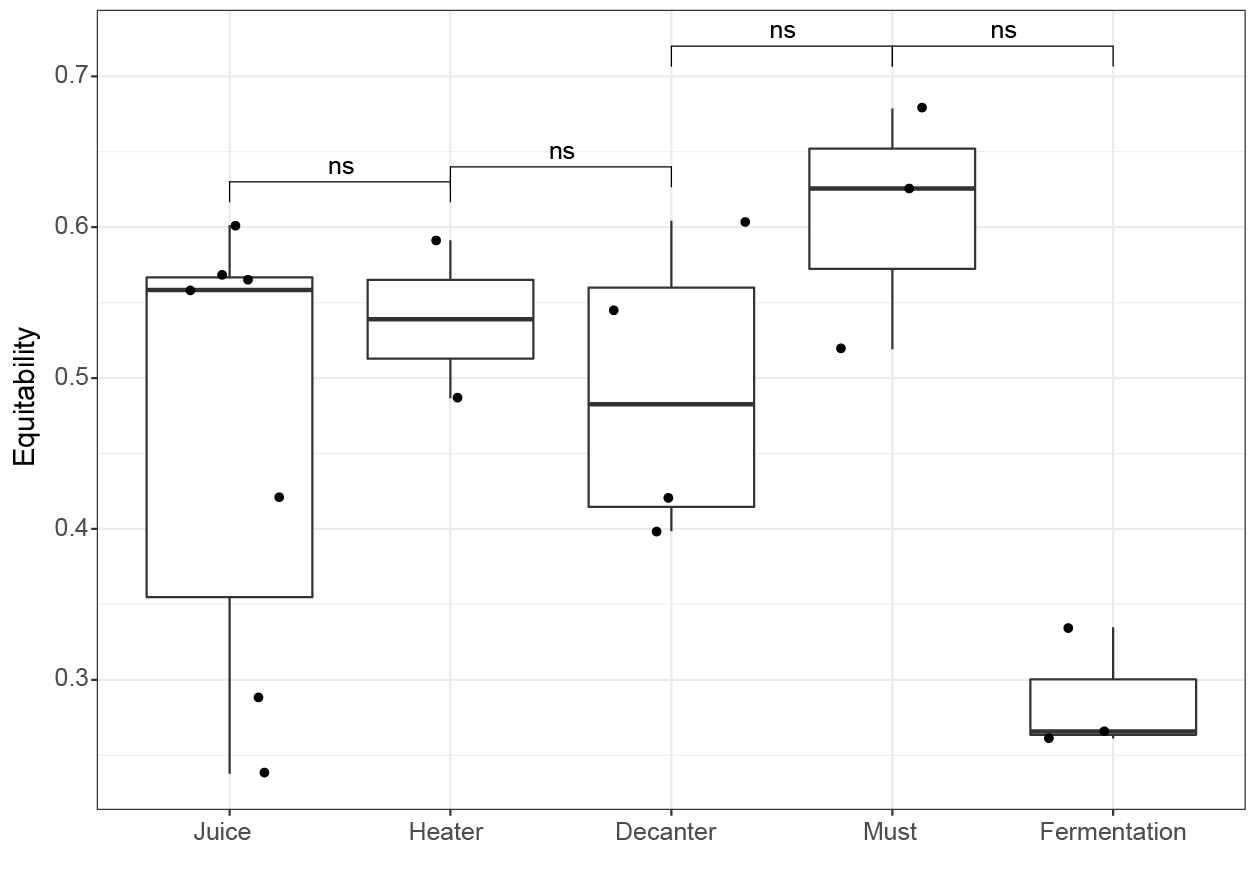

**Supplementary figure 3:** Equitability index in each step of ethanol production.

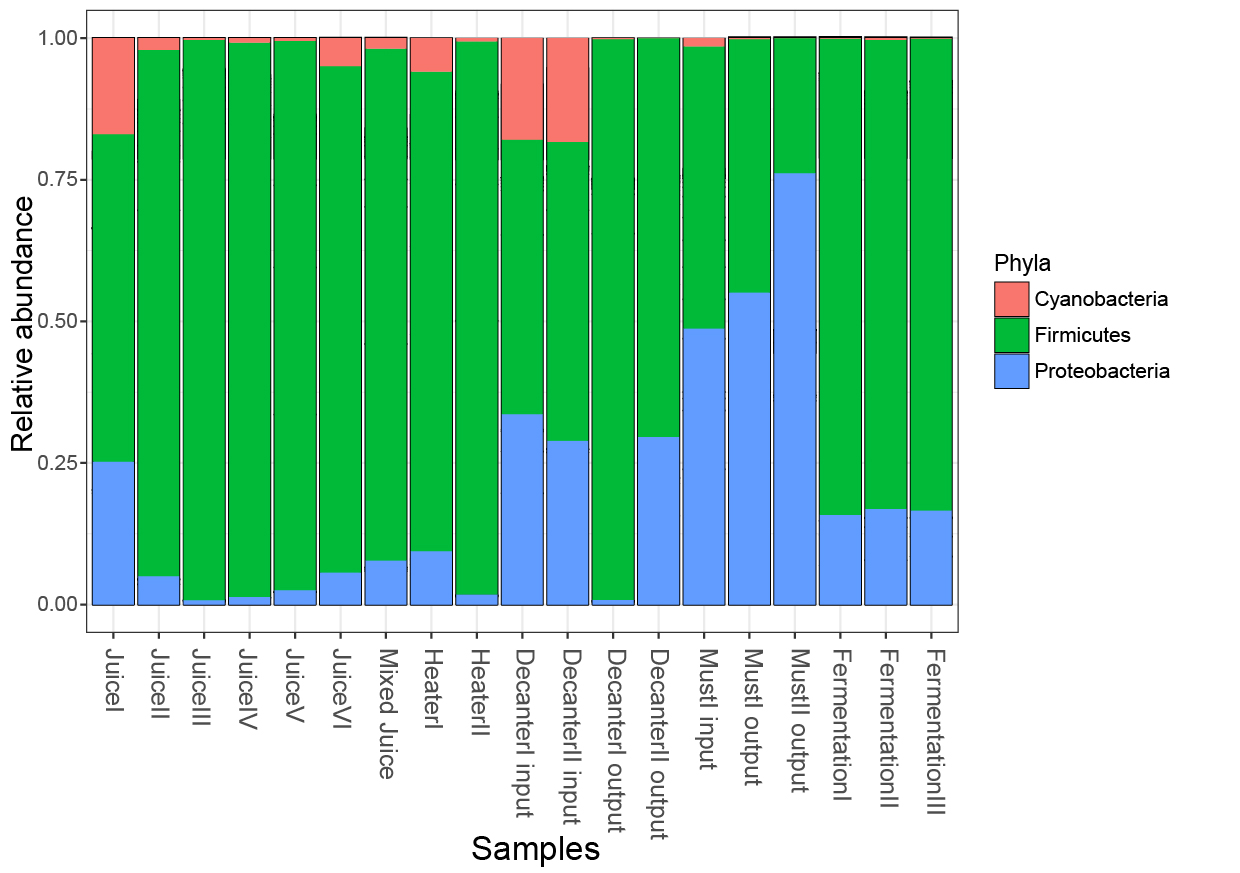

**Supplementary figure 4:** Relative abundance of the main phyla found in each step of ethanol production.

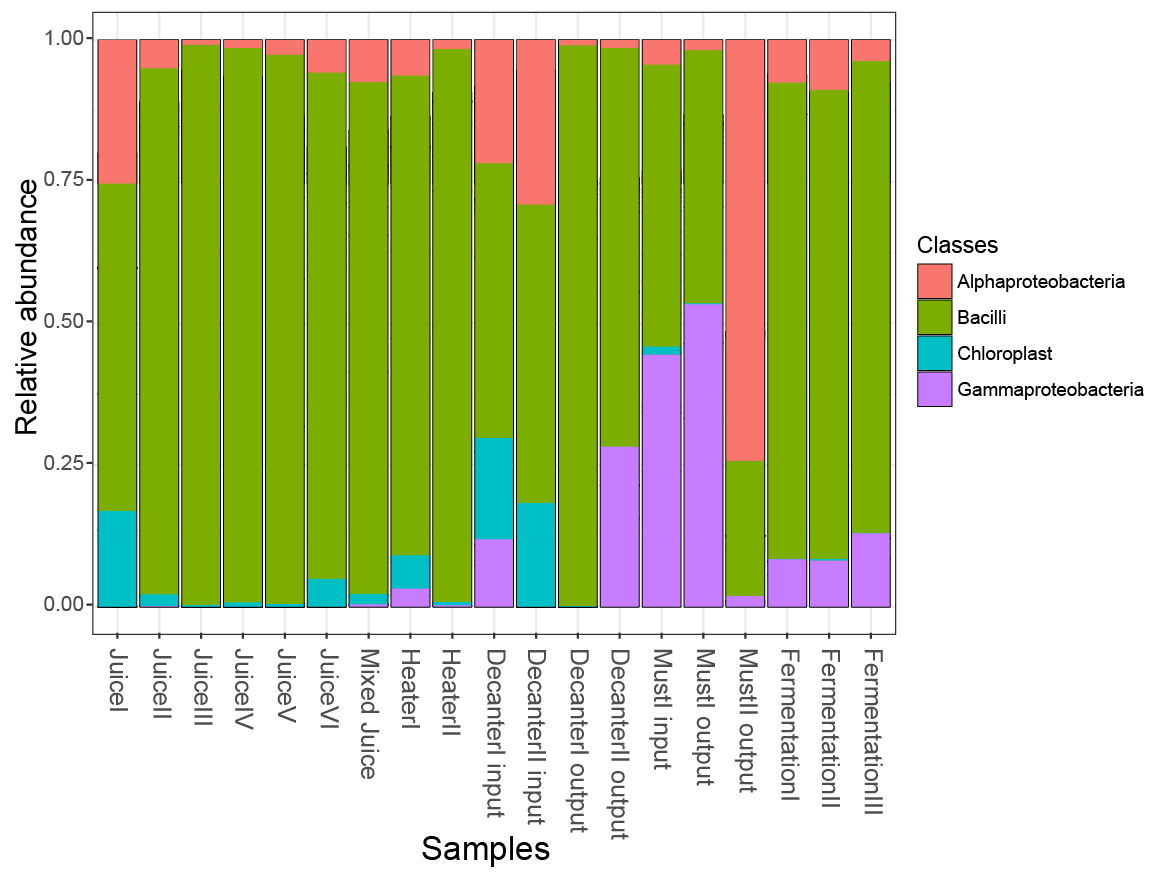

**Supplementary figure 5:** Relative abundance of the main classes found in each step of ethanol production.

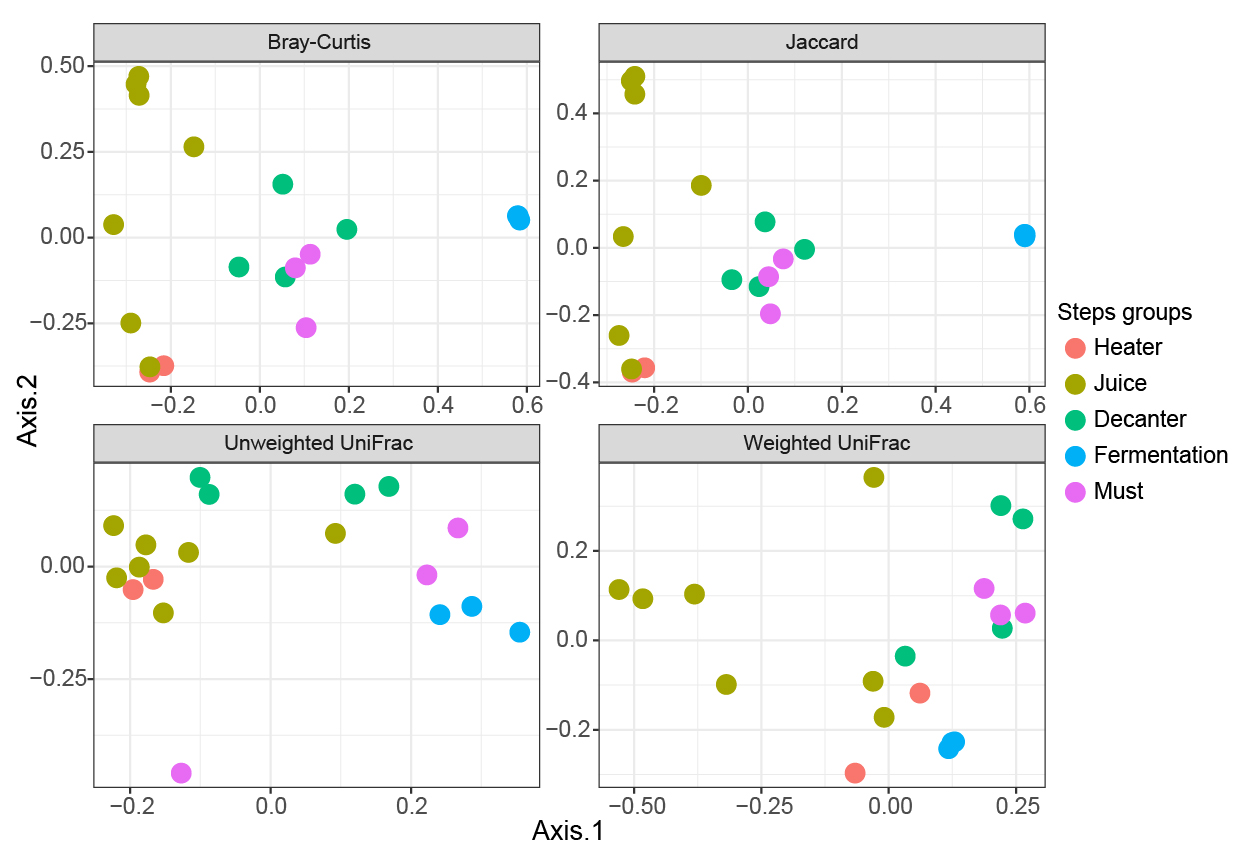

**Supplementary figure 6:** Principal Coordinates Analysis (PCoA) using Bray-Curtis, Jaccard, Unweighted and Weighted UniFrac dissimilarity matrix for samples of ethanol production process.
